# Evolution of mechanisms that control mating in *Drosophila* males

**DOI:** 10.1101/177337

**Authors:** Osama M. Ahmed, Aram Avila-Herrera, Khin May Tun, Paula H. Serpa, Justin Peng, Srinivas Parthasarathy, Jon-Michael Knapp, David L. Stern, Graeme W. Davis, Katherine S. Pollard, Nirao M. Shah

## Abstract

Genetically wired neural mechanisms inhibit mating between species because even naive animals rarely mate with other species. These mechanisms can evolve through changes in expression or function of key genes in specific sensory pathways or central circuits. Gr32a is a gustatory chemoreceptor that, in *D. melanogaster*, is essential to inhibit interspecies courtship and sense quinine. Similar to *D. melanogaster*, *D. simulans* Gr32a is expressed in foreleg tarsi, sensorimotor appendages that inhibit interspecies courtship in both species, and it is required to sense quinine. Nevertheless, Gr32a is not required to inhibit interspecies mating by *D. simulans* males. However, and similar to its function in *D. melanogaster*, Ppk25, a member of the Pickpocket family, promotes conspecific courtship in *D. simulans*. Taken together, we have identified shared as well as distinct evolutionary solutions to chemosensory processing of tastants as well as cues that inhibit or promote courtship in two closely related *Drosophila* species.

## INTRODUCTION

A species can be defined operationally as a set of organisms that share a gene pool and successfully breed with each other (Darwin, 1860; Dobzhansky, 1937; Mayr, 1988). An important tenet of evolutionary theory holds that species rarely interbreed, thereby preserving the advantages conferred by allele combinations that occur in conspecific gene pools (Mayr, 1988; Mayr and Dobzhansky, 1945; Orr, 2005; Orr et al., 2004). This immediately suggests that mechanisms that preclude interbreeding must evolve rapidly and facilitate reproductive isolation between species (Coyne and Orr, 1989; Mendelson, 2003). Given that even individuals from closely related species rarely attempt to mate with each other, the genetically-wired neural pathways underlying behavioral barriers to interbreeding must evolve rapidly. How such neural pathways evolve is poorly understood.

Drosophilid species provide a facile model to understand how neural pathways that inhibit interspecies courtship have evolved. There are ∼1,500 documented drosophilid species, many of which co-exist in overlapping habitats (Jezovit et al., 2017; Markow, 2015). They engage in well-described, species-typical stereotyped courtship rituals, and many components of the genetic and neural networks that regulate courtship of *D. melanogaster*, the most intensively studied drosophilid, are well defined (Bastock and Manning, 1955; Clowney et al., 2015; Demir and Dickson, 2005; Gill, 1963; Greenspan and Ferveur, 2000; Hall, 1978, 1994; Hotta and Benzer, 1976; Kallman et al., 2015; Kohatsu et al., 2011; Lin et al., 2016; Manoli et al., 2005; Pavlou and Goodwin, 2013; Ryner et al., 1996; Spieth, 1952; Thistle et al., 2012; Tootoonian et al., 2012). Moreover, we have previously demonstrated that sensory neurons expressing the gustatory chemoreceptor Gr32a are necessary to suppress interspecies courtship by *D. melanogaster* males (Fan et al., 2013). In addition, Gr32a is required to recognize cuticular hydrocarbons on non-*melanogaster* drosophilids and to inhibit interspecies mating. Strikingly, Gr32a is also necessary to inhibit courtship displays toward the closely related *D. simulans* that last shared an ancestor with *D. melanogaster* ∼3-5 million years ago (mya) (David et al., 2007; Tamura et al., 2004). *D. simulans* and *D. melanogaster* are cosmopolitan species that co-exist in shared habitats around the world {reviewed in (Jezovit et al., 2017)}. These two species are very similar in appearance and were only identified as distinct species upon close examination of male genitalia and the observation that they rarely courted each other (Sturtevant, 1919, 1920). Here we have examined how the Gr32a chemosensory pathway has evolved to inhibit interspecies courtship in *D. simulans*.

## RESULTS

### The chemosensory pathway that inhibits interspecies courtship is conserved

*D. melanogaster* males exhibit a highly ritualized courtship sequence such that, having oriented head first to a potential mate, they tap it with their foreleg tarsi. This tapping restricts subsequent steps of courtship to conspecific individuals because *D. melanogaster* males lacking foreleg tarsi continue to court conspecifics as well as individuals from other drosophilid species (Fan et al., 2013; Manning, 1959). *D. simulans* males also tap potential mates with their foreleg tarsi early in courtship, and we therefore tested whether these sensorimotor appendages serve a similar function in *D. simulans* males (*Figure 1A*). We surgically ablated foreleg tarsi in adult *D. simulans* males and tested them for courtship toward conspecifics and individuals of other species. We observed that male *D. simulans* lacking foreleg tarsi courted closely (*D. melanogaster*) and distantly (*D. virilis*, shared last common ancestor ∼40 mya) related drosophilids as well as male conspecifics (*Figure 1B–E* and *Figure 1–figure supplement 1A*). Importantly, *D. simulans* males lacking foreleg tarsi, like their *D. melanogaster* counterparts (Fan et al., 2013), also courted conspecific females (*Figure 1B,C*). Both *D. melanogaster* and *D. simulans* tarsiless males exhibited reduced courtship toward conspecific females compared to sham-manipulated males. This may reflect a diminution in their ability to pursue the female effectively because of impaired locomotion or loss of neurons that detect attractive pheromones and promote courtship. Regardless, the behavior of tarsiless *D. simulans* males was in contrast to intact males who exhibited very low levels of courtship toward other species and conspecific males.

**Figure 1:**
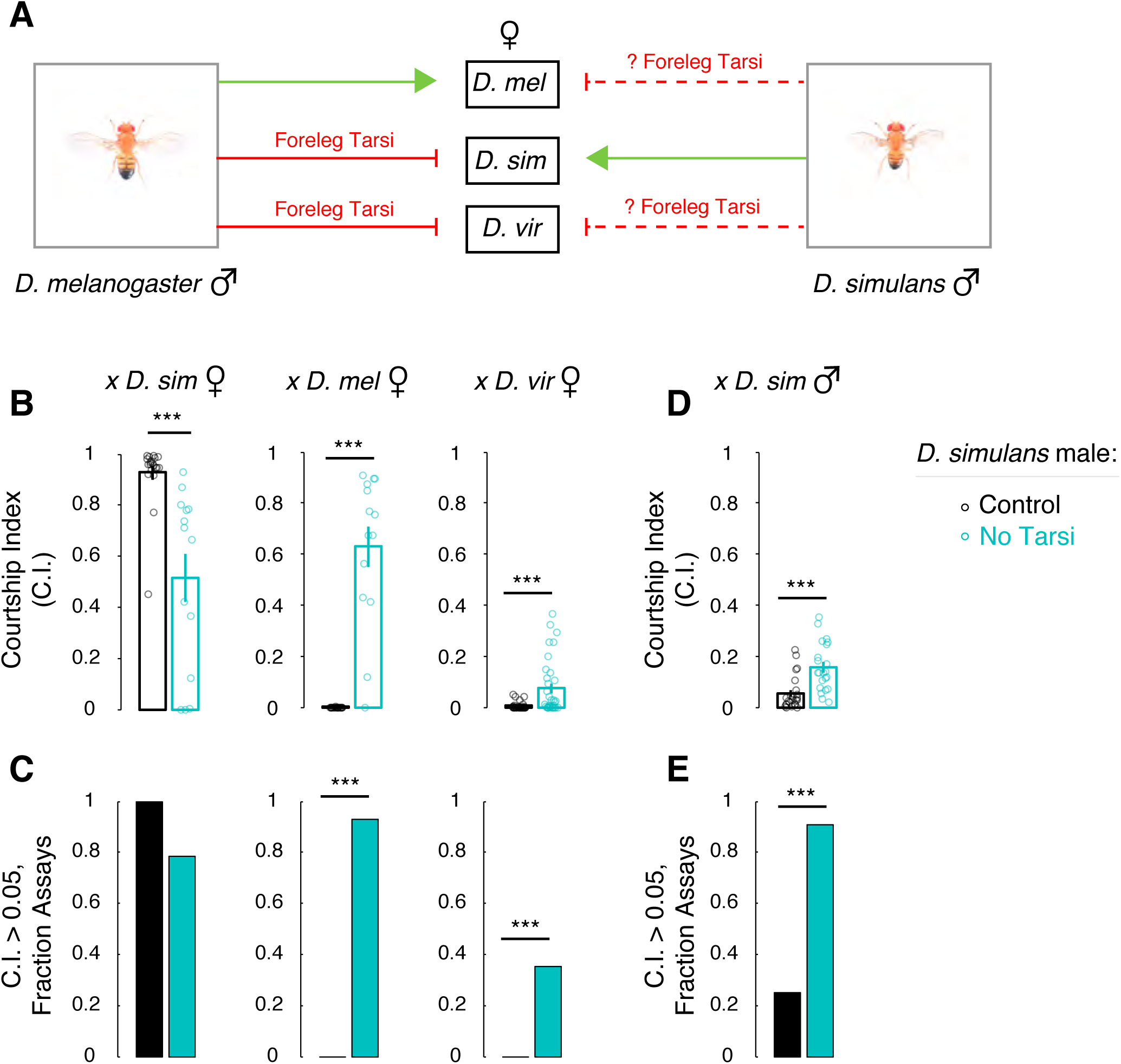
*D. simulans* male foreleg tarsi inhibit courtship of other species and are not essential for courtship of conspecific females. (A) We tested whether, similar to *D. melanogaster* males, foreleg tarsi also inhibited interspecies courtship by *D. simulans* males. (B) *D. simulans* males lacking foreleg tarsi court conspecific, *D. melanogaster*, and *D. virilis* females. (C) *D. simulans* males lacking foreleg tarsi are more likely to show intense courtship toward *D. melanogaster* and *D. virilis* females. (D) *D. simulans* males lacking foreleg tarsi show more courtship toward conspecific males. (E) *D. simulans* males lacking foreleg tarsi are more likely to show intense courtship toward conspecific males. Mean ± SEM; CI = fraction time spent courting target fly; each circle denotes CI of one male; n = 14 - 41/cohort; ***p<0.001. See also *Figure 1–figure supplement 1*.

The hydrocarbon 7-tricosene is present on the cuticle of *D. simulans* but not *D. melanogaster* females, and it functions as an aphrodisiac and a repulsive chemosensory cue for males of these two species, respectively (Billeter et al., 2009; Fan et al., 2013; Ferveur, 2005; Lacaille et al., 2007; Wang et al., 2011). Consistent with this notion, WT *D. simulans* courted *D. yakuba* females, whose cuticle is enriched in 7-tricosene, albeit with significantly lower intensity compared to conspecific females (p-value < 0.001; n = 20 - 22 males/cohort; see *Figure 1–figure supplement 1B,C* and *Figure 1B*). Removal of foreleg tarsi in the males did not further increase courtship toward *D. yakuba* females (*Figure 1–figure supplement 1B,C*). Our findings therefore suggest that multiple pathways exist in *D. simulans* to inhibit interspecies courtship toward other drosophilids. Nevertheless, and similar to *D. melanogaster* males, severing foreleg tarsi of *D. simulans* males disinhibits courtship toward multiple reproductively futile targets without abolishing courtship with conspecific females.

### Gr32a expression is conserved in *D. simulans* foreleg tarsi

Foreleg tarsi contain chemosensory neurons that presumably detect contact-dependent cues during tapping such that detection of non-conspecific cues inhibits subsequent steps in courtship. Indeed, the chemoreceptor-encoding gene *Gr32a* is expressed in specific sensory neurons in distal foreleg tarsi of *D. melanogaster* (Koganezawa et al., 2010; Miyamoto and Amrein, 2008; Moon et al., 2009; Scott et al., 2001; Thistle et al., 2012; Thorne et al., 2004), and it is required to inhibit interspecies courtship (Fan et al., 2013) (*Figure 2A*). Moreover, *Gr32a*-expressing neurons are functionally necessary and sufficient to inhibit interspecies courtship displays. Given that the genome of *D. simulans* also encodes an ortholog of *Gr32a* (*Drosophila* 12 Genomes Consortium et al., 2007), we wondered whether this gene is expressed in foreleg tarsi of this species. The Gr gene family members are transcribed at very low levels (Clyne et al., 2000; Dunipace et al., 2001; Moon et al., 2009; Scott et al., 2001), and previous efforts have utilized cognate upstream promoter regions to visualize expression of Gr32a using the GAL4/UAS system (Park and Kwon, 2011; Scott et al., 2001; Wang et al., 2004; Weiss et al., 2011). These studies have shown that ∼3.8 kb of *D. melanogaster* genomic DNA immediately upstream of the start codon is sufficient to drive reporter expression in subsets of neurons in chemosensory organs known to express Gr32a (Scott et al., 2001; Wang et al., 2004). Similar stretches of genomic DNA immediately upstream of the start codon are also sufficient to drive reporter expression of other Grs in various chemosensory neurons (Weiss et al., 2011), indicating a conserved regulatory logic of expression for this gene family in *D. melanogaster*. Accordingly, we subcloned ∼3.8 kb of genomic DNA upstream of the *D. simulans* Gr32a start codon and used it to drive GAL4 expression (Gr32a^sim^-GAL4) in transgenic *D. simulans* and *D. melanogaster* flies (Pfeiffer et al., 2010; Stern et al., 2017) (*Figure 2B*). Transgene expression was visualized in progeny bearing this allele as well as the fluorescent protein Citrine under control of UAS promoter sequence (Inagaki et al., 2014) (*Figure 2C,D*). We observed Citrine expression in a small subset of neurons in distal tarsal segments (T4, T5) of *D. simulans* and *D. melanogaster*, demonstrating that regulatory sequences in the *D. simulans Gr32a* locus drive reporter expression in foreleg tarsi of both species (*Figure 2C,D,G*).

**Figure 2:**
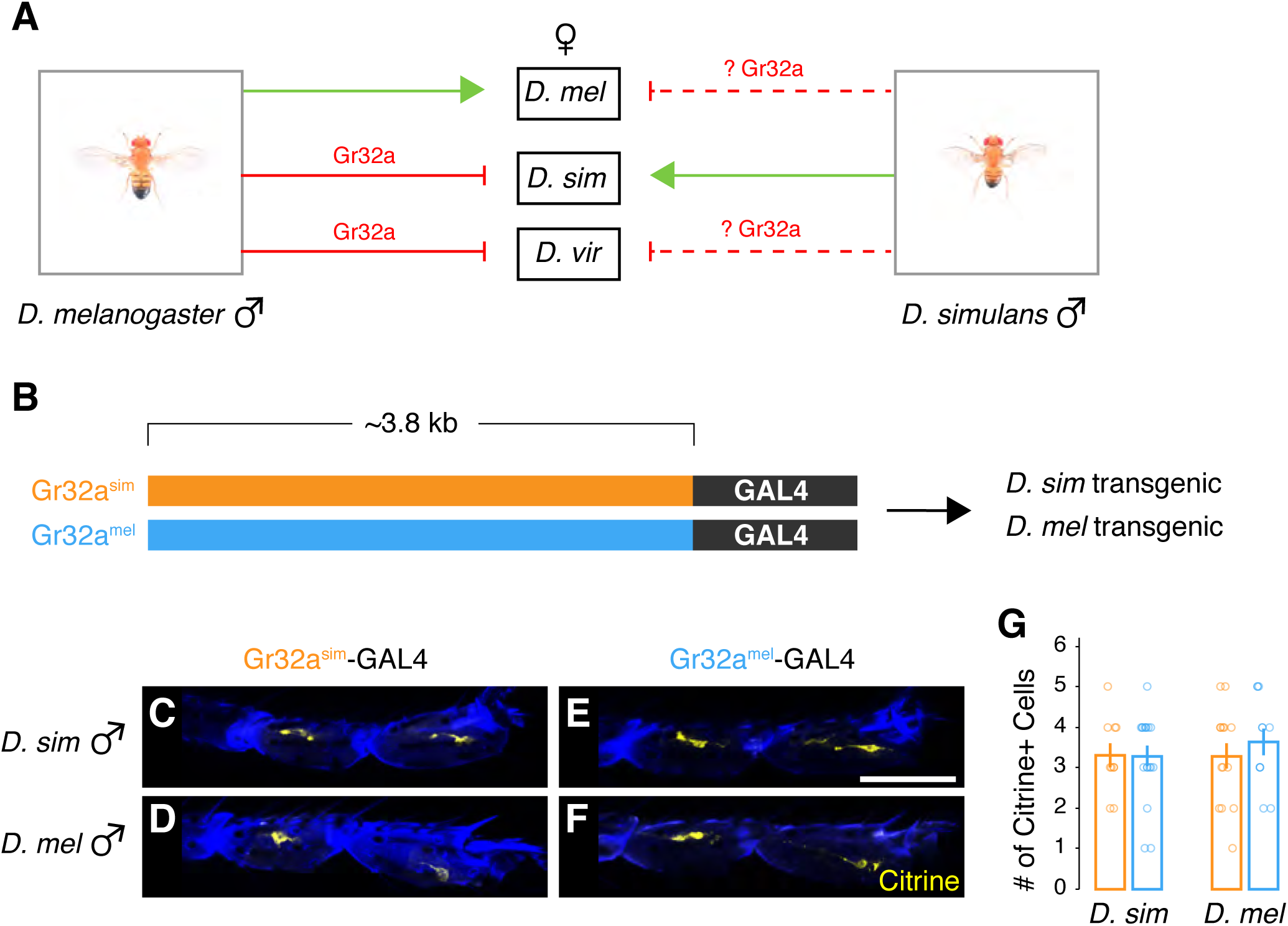
A regulatory region in the *Gr32a* locus is functionally conserved. (A) We sought to determine whether, similar to *D. melanogaster*, Gr32a was expressed in *D. simulans* foreleg tarsi. (B) Schematic of transgenic constructs using a DNA sequence 5’ of Gr32a start codon from *D. simulans* (orange) and *D. melanogaster* (blue) to drive GAL4 expression. (C-F) Gr32a^sim^-GAL4 and Gr32a^mel^-GAL4 each drive comparable citrine expression in distal tarsal segments T4 and T5 in both *D. simulans* and *D. melanogaster* male forelegs. (G) Quantification of data shown in histological panels (C-F). Mean ± SEM; each circle denotes number of Citrine+ cells per male foreleg tarsi per; n = 11 - 18/genotype; scale bar = 50 μm. See also *Supplementary Table 1* and *Figure 2–figure supplement 1*.

We next wanted to test whether the ∼3.8 kb regulatory DNA sequence from these two species drives expression in the same tarsal neurons. We therefore generated *D. melanogaster* flies harboring GAL4 under control of conspecific ∼3.8 kb DNA sequence 5’ of Gr32a such that this transgene (Gr32a^mel^-GAL4) was inserted into the same landing site that we had utilized for Gr32a^sim^-GAL4 (*Figure 2B,D,F*). This strategy ensured that transgenes from the two species were regulated by the identical genomic context flanking the insertion site. Importantly, Gr32a^mel^-GAL4 regulated reporter expression in *D. melanogaster* foreleg tarsi (T4, T5) as described previously for other GAL4 alleles of Gr32a (Fan et al., 2013; Miyamoto and Amrein, 2008; Moon et al., 2009; Scott et al., 2001). In *D. melanogaster* flies bearing both Gr32a^mel^-GAL4 and Gr32a^sim^-GAL4, we observed a similar number of Citrine+ foreleg tarsal neurons compared to flies bearing each of these GAL4 drivers individually (*Figure 2–figure supplement 1A*). Together, these data are consistent with the notion that the upstream regulatory region of Gr32a in the two species is functionally conserved and sufficient to drive expression in the same foreleg tarsi neurons of *D. melanogaster*.

To investigate whether these regulatory features also function in *D. simulans*, we tested whether the ∼3.8 kb genomic DNA upstream of *D. melanogaster* Gr32a would drive expression in foreleg tarsal neurons of *D. simulans*. Accordingly, we inserted Gr32a^mel^-GAL4 into the identical landing site we used to generate *D. simulans* flies bearing Gr32a^sim^-GAL4 (*Figure 2B,E*). As before, transgene expression was visualized in progeny bearing this allele and Citrine under control of UAS. We observed Citrine expression in 3-4 neurons restricted to distal tarsal segments (T4, T5) of *D. simulans* in a pattern mirroring that observed in *D. simulans* bearing Gr32a^sim^-GAL4 (*Figure 2C,E,G*). Given that all transgenes we built in *D. simulans* were inserted into a single landing site that has afforded us reliable and non-leaky expression, we cannot directly test whether the same neurons were labeled by Gr32a^sim^-GAL4 and Gr32a^mel^-GAL4 in this species. Nevertheless, our findings strongly suggest that similar *cis* and *trans* regulatory features regulate Gr32a expression in foreleg tarsi of the two species.

In fact, we find that the ∼3.8 kb stretch of genomic DNA appears conserved across multiple insect species (mean nucleotide conservation phyloP score = 1.4; see *Figure 2–figure supplement 1B–D*). Coding exons for another gene (*D. melanogaster CG6201*) contribute to this sequence similarity, but some of the most conserved blocks of sequence are intergenic regions directly upstream of the Gr32a start codon and 5’ UTR (*Figure 2–figure supplement 1C*). Overall, nucleotide substitutions have occurred in this region at 42.5% the rate of 4-fold degenerate sites in protein-coding exons, significantly slower than expected under a neutral model of DNA evolution for these insects (p < 1 x 10^−5^, see Materials and methods for details; *Figure 2–figure supplement 1D*). Within *D. melanogaster* and *D. simulans*, >95% of the DNA sequence is identical across this ∼3.8 kb region (*Figure 2–figure supplement 1B*). To examine sequence differences at single nucleotide resolution, we tested each position for a faster or slower rate of DNA substitutions in *D. melanogaster* than expected, given the rate in *D. simulans* and the other 25 insects. We also conducted the comparable test for *D. simulans*. This analysis revealed that very few bases in the ∼3.8 kb region are evolving faster than expected in *D. melanogaster* or *D. simulans* (>99% bases with phyloP score > −2; *Figure 2–figure supplement 1E,F*). Because the ∼3.8 kb region is highly conserved across diverse insects and *D. melanogaster* and *D. simulans* diverged from a common ancestor very recently, it was difficult to detect whether this stretch of DNA is evolving slower than expected subsequent to speciation from this shared ancestor. Taken together, our findings show that this ∼3.8 kb region is conserved in sequence and function in *D. melanogaster* and *D. simulans* such that it is sufficient to drive expression in neurons of foreleg tarsi, a structure that inhibits interspecies courtship by males of the two species.

### Gr32a and Gr33a are not essential to inhibit interspecies courtship in *D. simulans* males

*D. melanogaster* males mutant for Gr32a exhibit increased courtship toward heterospecific females (Fan et al., 2013). We therefore tested whether Gr32a was essential to inhibit interspecies courtship in *D. simulans* males (*Figure 3A*). We used two guide RNAs targeting distinct sequences in the first coding exon of *D. simulans* Gr32a to generate three different mutant alleles via the CRISPR/Cas9 system (Bassett and Liu, 2014; Bassett et al., 2013; Gokcezade et al., 2014) (*Figure 3–figure supplement 1A–C*). Two of the alleles (*Gr32a*^*Δ10*^ and *Gr32a*^*Δ26*^) are predicted to lead to 10 and 26 bp deletions in the first coding exon that result in a frame-shift and premature stop codon; these likely encode a non-functional Gr32a chemoreceptor protein (*Figure 3–figure supplement 1C,D*). The third allele (*Gr32a*^*Δ141*^) has a 141 bp deletion that is predicted to eliminate 47 amino acids from the predicted N-terminal intracellular domain of this chemoreceptor (*Figure 3–figure supplement 1B–D* and *Figure 3– figure supplement 2*). We back-crossed these mutant alleles into a WT background ≥5 times to remove any potential off-target mutations and only then subjected resulting progeny to testing in behavioral and other assays. We also confirmed that these deletions were present within the mRNAs transcribed from each of the three alleles *in vivo* in adult flies (*Figure 3–figure supplement 1B*). We next tested *D. simulans* males homozygous mutant for these three *Gr32a* alleles for courtship displays toward conspecifics and members of other species. We observed that each of these three mutants courted conspecific females at levels indistinguishable from WT controls (*Figure 3B,C*). Moreover, and unlike *D. melanogaster* males mutant for Gr32a, these mutants did not show a significant increase in courtship toward conspecific males or *D. melanogaster, D. yakuba,* or *D. virilis* females (*Figure 3B–E* and *Figure 3–figure supplement 1E,F*) (Fan et al., 2013; Miyamoto and Amrein, 2008). Our findings therefore indicate a divergence in behavioral function of Gr32a between *D. simulans* and *D. melanogaster*. This is consistent with sequence analyses of Gr family members that suggest that bitter-sensing Grs such as Gr32a may be evolving rapidly (Gardiner et al., 2009; McBride et al., 2007). In summary, Gr32a mutant *D. simulans* males do not show elevated courtship toward other species, a finding in sharp contrast to Gr32a null *D. melanogaster* males who court other species avidly (Fan et al., 2013).

**Figure 3:**
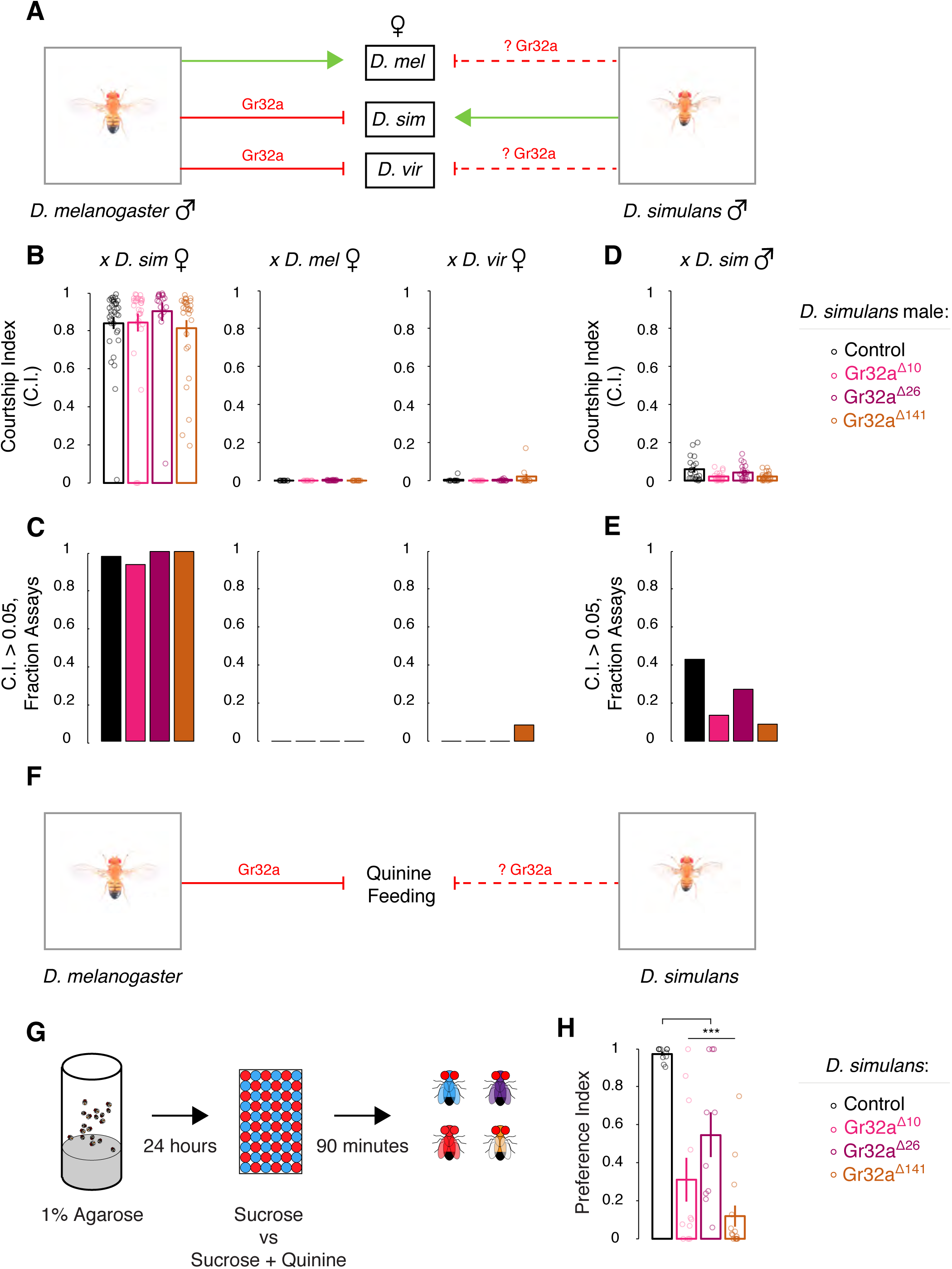
Gr32a is not required to inhibit interspecies courtship but is essential for quinine sensing in *D. simulans*. (A) We tested whether, similar to *D. melanogaster* males, Gr32a inhibits interspecies courtship by *D. simulans* males. (B, C) WT and Gr32a mutant *D. simulans* males court conspecific but not *D. melanogaster* or *D. virilis* females. (D, E) WT and Gr32a mutant *D. simulans* males show similar low levels of courtship toward conspecific males. (F) We tested whether, similar to *D. melanogaster*, Gr32a inhibits feeding on quinine-containing food in *D. simulans*. (G) Schematic of feeding assay for starved *D. simulans* given choice of colored food containing sucrose or sucrose and quinine. Flies with blue, red, purple, or no food dye colored abdomens were enumerated after exposure to food for 90 min. (H) Significant decrease in preference by *Gr32a* mutant *D. simulans* for food containing only sucrose. Mean ± SEM; (B-E) each circle denotes CI of one male, and n = 11 - 34/genotype. (G, H) Preference Index = {(# flies that ate sucrose-only food + 0.5*(purple flies)}/(# flies that ate); each circle denotes Preference Index for one experiment; 106 ± 6 *D. simulans* of each genotype were used/experiment; n = 11 - 15 experiments/genotype; ***p<0.001. Please see *Supplementary Tables 1-3* and *Figure 3–figure supplements 1-5*.

Gr33a is co-expressed with Gr32a in foreleg tarsi in *D. melanogaster*, and it is required to inhibit intermale but not interspecies courtship in males of this species (Fan et al., 2013; Moon et al., 2009). Gr33a is also encoded in the *D. simulans* genome (Drosophila 12 Genomes Consortium et al., 2007), and we wondered if this chemoreceptor had evolved to inhibit interspecies courtship in this species. We tested this hypothesis by using CRISPR to generate two distinct alleles of *Gr33a*, one with a 10 bp deletion (*Gr33a*^*Δ10*^) that leads to a frame-shift and premature stop codon and the other encompassing an in-frame deletion (96 bp, *Gr33a*^*Δ96*^) (*Figure 3–figure supplement 3A–D*). As before, we also back-crossed these mutants ≥5 times into a WT background to remove potential mutations resulting from off-target events. We confirmed the presence of these deletions within endogenously transcribed message in adult flies (*Figure 3–figure supplement 3B*). In behavioral testing, male *D. simulans* mutant for each of these alleles courted conspecific females similar to WT controls, but they did not display significant courtship toward conspecific males or *D. melanogaster, D. yakuba,* or *D. virilis* females (*Figure 3–figure supplement 3E–H* and *Figure 3–figure supplement 4*). Taken together, our results indicate that chemosensory receptor-mediated inhibition of courtship toward reproductively futile targets – conspecific males and members of other species – has diverged between the closely related *D. melanogaster* and *D. simulans*.

### Both Gr32a and Gr33a are required functionally in *D. simulans* to detect the aversive tastant quinine

In addition to inhibiting courtship of reproductively futile targets by *D. melanogaster* males, Gr32a and Gr33a are essential for chemosensory neuronal responses to quinine as well as a behavioral aversion to this bitter tastant (Lee et al., 2010; Moon et al., 2009). Previous studies have shown that chemoreceptors can evolve to enable food-sensing in different species inhabiting particular ecological niches (Baldwin et al., 2014; Jordt and Julius, 2002; Prieto-Godino et al., 2017; Wisotsky et al., 2011). We therefore wondered if Gr32a and Gr33a were required in *D. simulans* for the aversive response to quinine (*Figure 3F* and *Figure 3–figure supplement 5A*). We tested this by performing a feeding preference assay in which starved flies were offered the choice to feed on food containing low concentration of sugar (1 mM sucrose) or high concentration of sugar (5 mM sucrose) spiked with quinine (0.5 mM) (Montell, 2009; Tanimura et al., 1982) (*Figure 3G*). In contrast to WT *D. simulans* that strongly preferred feeding on the low concentration of sugar, flies mutant for either Gr32a or Gr33a showed a highly significant reduction in preference for feeding on sugar alone (*Figure 3H* and *Figure 3– figure supplement 5B*). In summary, Gr32a and Gr33a are essential for sensing quinine in *D. simulans*. We cannot exclude the possibility that all five Gr32a and Gr33a mutations we have generated in *D. simulans* represent alleles that disrupt sensing quinine but not chemosensory cues from other species. This possibility could be tested when deficiencies spanning Gr32a and Gr33a become available in this species. In any event, our present findings demonstrate that the function of Gr32a and Gr33a in sensing quinine is conserved between *D. melanogaster* and *D. simulans*.

### Ppk25 promotes conspecific courtship in *D. simulans* males

Our findings show that chemosensory receptor mechanisms that inhibit courtship of reproductively futile targets in *D. melanogaster* are not employed in *D. simulans*. We wondered whether genetic loci that promote courtship had also differentiated between these two species. Many distinct loci have previously been shown to promote courtship of *D. melanogaster* males toward conspecific females {reviewed in (Dickson, 2008; Yamamoto and Koganezawa, 2013)}. We chose to test the function of the Ppk25 pickpocket ion channel subunit that is expressed in foreleg tarsi chemosensory neurons and appears to exclusively promote courtship in *D. melanogaster* (*Figure 4A*) (Clowney et al., 2015; Kallman et al., 2015; Lin et al., 2005; Starostina et al., 2012; Vijayan et al., 2014). We generated two distinct alleles of Ppk25 in *D. simulans* via CRISPR/Cas9, one of which has a 2 bp insertion and the other has a 4 bp deletion in the first coding exon (*Figure 4–figure supplement 1A-C*). Both alleles are predicted to lead to frame-shifts and premature stop codons, and are likely therefore to encode null alleles of this gene (*Figure 4–figure supplement 1D*). Subsequent to ≥5 back-crosses to minimize CRISPR-generated off-target mutations in the background, we tested males homozygous mutant for each allele in courtship assays. *D. melanogaster* males, similar to *D. simulans* males, use multiple cues to initiate courtship with conspecifics, and *D. melanogaster* Ppk25 is required for male courtship in the dark (Boll and Noll, 2002; Jezovit et al., 2017; Kohatsu and Yamamoto, 2015; Krstic et al., 2009; Lin et al., 2005; Spieth, 1974). Unlike *D. melanogaster*, *D. simulans* males exhibit high levels of conspecific courtship only under bright illumination (Grossfield, 1971; Jezovit et al., 2017) (*Figure 4–figure supplement 2*). We therefore tested whether Ppk25 modulated courtship by *D. simulans* males in bright light or dark conditions. In keeping with a role for this gene in promoting courtship, we observed that *D. simulans* males mutant for Ppk25 showed reduced courtship of conspecific females compared to WT control males in the dark (*Figure 4B,C*). These mutants also showed subtle, but significant, reduction in courtship under bright illumination, suggesting a more stringent requirement for Ppk25 in promoting this behavior (*Figure 4D,E*). Similar to WT controls, *D. simulans* males mutant for Ppk25 did not display elevated courtship to other drosophilids (*Figure 4–figure supplement 3*), indicating that it does not function in this species to inhibit interspecies courtship. In fact, we found that Ppk25 mutant *D. simulans* had subtle, but significant reduction, in the low levels of courtship normally displayed by WT males to *D. yakuba* females (*Figure 4–figure supplement 3*). In summary, Ppk25 functions in both *D. melanogaster* and *D. simulans* to promote WT courtship displays.

**Figure 4:**
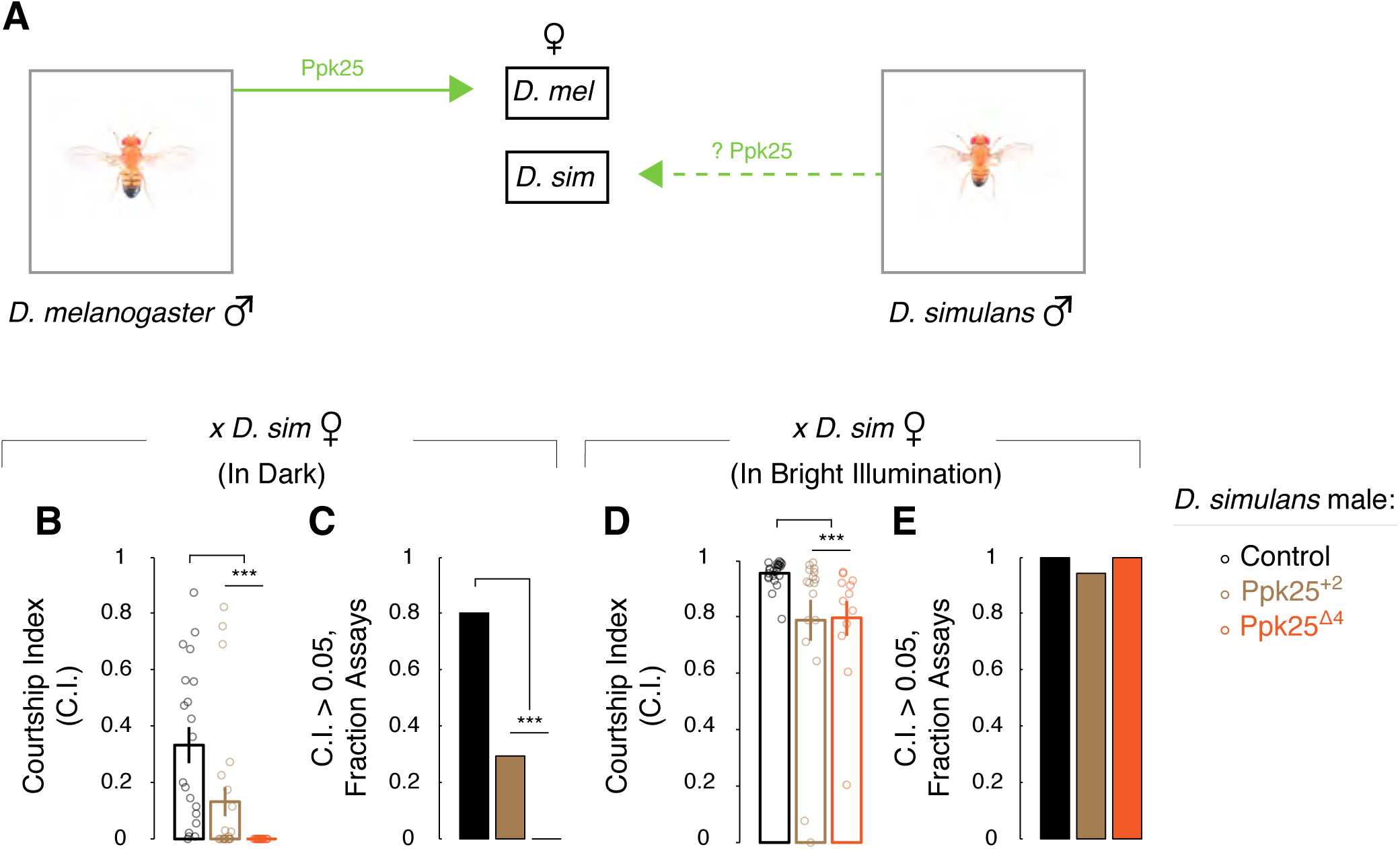
Ppk25 promotes conspecific courtship by *D. simulans* males. (A) We tested whether, similar to *D. melanogaster*, Ppk25 promotes conspecific courtship by *D. simulans* males. (B-D) *Ppk25* mutant *D. simulans* males show decreased courtship toward conspecific females. (E) No difference between WT and *Ppk25* mutant *D. simulans* males in percent assays with high levels of courtship of conspecific females. Mean ± SEM; each circle denotes CI for one male; n = 12 - 24/genotype; ***p<0.001. Please see *Supplementary Tables 1,3* and *Figure 4–figure supplements 1,2*.

## DISCUSSION

The study of changes in morphological or other traits across evolution continues to be vigorously investigated (Carroll, 2008). We have examined whether a pathway that inhibits interspecies courtship in *D. melanogaster* might perform a similar function in *D. simulans*, a sympatric species that diverged recently. Similar to *D. melanogaster*, we find that male foreleg tarsi in *D. simulans* are essential to inhibit this behavior but not for conspecific courtship. Gr32a, a chemoreceptor that is expressed in *D. melanogaster* foreleg tarsi neurons and required to inhibit interspecies courtship, is also expressed in foreleg tarsi neurons of *D. simulans*. Moreover, regulatory DNA sequence elements within the *Gr32a* locus of these species are likely functionally conserved as they appear sufficient to confer reporter expression in foreleg tarsi neurons of the cognate as well as the heterologous species. Remarkably however, Gr32a is not essential for inhibiting interspecies courtship in *D. simulans*. Thus, the male forelegs of both species inhibit interspecies courtship but chemoreceptor control of this behavior has diverged within ∼3-5 million years.

Gr32a may not be essential to suppress interspecies courtship in *D. simulans*, but it is possible that Gr32a neurons in foreleg tarsi still function to inhibit this behavior. Accordingly, we wished to functionally silence Gr32a-expressing neurons in an effort to determine if these neurons inhibit interspecies courtship by *D. simulans* males. However, it was technically challenging to generate the reagents required for these experiments (Kir2.1, tetanus toxin light chain, *shibire*^*ts*^) (Luo et al., 2008) in this species, despite numerous attempts. *D. simulans* males sense aversive cues on the cuticle of *D. melanogaster* females, suggesting that they use a chemosensory pathway to avoid mating with other species (Billeter et al., 2009). Given that Gr32a does not serve this function in *D. simulans*, what chemoreceptors might be employed to detect repellents that preclude interspecies courtship in this species? We find that the related chemoreceptor Gr33a, which is co-expressed in many Gr32a neurons in *D. melanogaster* foreleg tarsi, is not required to inhibit this behavior. It is possible that, in *D. simulans*, *Gr32a* and *Gr33a* function redundantly to inhibit interspecies courtship, a notion difficult to test directly because these loci are only 1 Mb apart in the genome. Moreover, even if this notion is correct, our findings still demonstrate a divergence in the function of Gr32a (and Gr33a) between *D. melanogaster* and *D. simulans*. In addition, the gustatory and ionotropic chemoreceptor families contain many members, and it is possible that a different chemoreceptor(s) functions to inhibit interspecies courtship by *D. simulans* males (Joseph and Carlson, 2015). Whether such a chemoreceptor functions in Gr32a-expressing or other neurons in foreleg tarsi also remains to be determined.

The divergence in chemoreceptor-mediated suppression of courtship between *D. melanogaster* and *D. simulans* does not reflect a global reorganization of molecular pathways that regulate courtship. We find that, similar to its role in *D. melanogaster*, Ppk25 is required to promote courtship toward conspecific females in *D. simulans*. Ppk25 is required to sense 7,11-heptacosadiene, an aphrodisiac cue, in *D. melanogaster* (Kallman et al., 2015; Starostina et al., 2012); however, 7,11-heptacosadiene is an aversive cue for *D. simulans* males such that they do not court targets coated with this pheromone (Billeter et al., 2009). Given these constraints, it will be interesting to understand how the chemosensory neurons expressing Ppk25 function in both species to promote conspecific courtship. In one scenario, Ppk25 does not directly sense such aphrodisiac pheromones, but rather it transduces activation of pheromone-sensing chemoreceptors that remain to be identified. If so, we anticipate that these neurons express different chemoreceptors between the species or that the chemoreceptor has evolved to sense species-specific aphrodisiacs.

Both Gr32a and Gr33a are required for avoidance of the bitter tastant quinine in *D. melanogaster* and *D. simulans*. Thus, the functions of Gr32a and Gr33a in avoiding quinine and inhibiting courtship of reproductively futile targets are evolutionarily dissociable between closely related species within a genus. Our findings demonstrate that although the same behavioral trait (tapping) and sensorimotor appendage (foreleg) inhibit courting of reproductively dead-end targets in *D. melanogaster* and *D. simulans*, the molecular mechanisms that preclude such courtship appear to have diverged between these species. Previous work from our and other labs shows that different genetic pathways control distinct quantitative aspects of behavioral subroutines (Ding et al., 2016; Greenwood et al., 2013; Weber et al., 2013; Xu et al., 2012). Together, these findings demonstrate that modifications in genetic pathways may be employed to gate a behavior or to implement quantitative changes in that behavior. We also find that although chemoreceptor mechanisms inhibiting interspecies courtship have differentiated between closely related species, a chemosensory pathway promoting courtship appears to have a similar positive valence in both species. It will be interesting to determine whether these courtship promoting and inhibiting pathways evolve in a similar pattern across other drosophilid species. Alternatively (Jacob, 1977; Luo, 2015), our findings may reflect the idiosyncratic nature of selective forces that exploit mutations in apparently random pathways to effect evolutionary change. It should be possible to distinguish between these interesting alternatives by studying mechanisms that regulate courtship in additional drosophilid species.

## MATERIALS AND METHODS

### Drosophila stocks

*D. simulans* (14021-0251.001), w^501^ *D. simulans* (14021-0251.195), *D. yakuba* (14021-0261.00), and *D. virilis* (15010-1051.00) were obtained from the Drosophila Species Stock Center at the University of California, San Diego. WT *D. melanogaster* were in the Canton-S background. *D. melanogaster* UAS-ReaChR::Citrine.VK05 was obtained from the Bloomington Drosophila Stock Center (#53749). Transgenic and CRISPR-mediated mutant flies were generated as described below.

### Generating *D. simulans Gr32a*, *Gr33a*, or *Ppk25* mutants

CRISPR guides were chosen from a list generated by flyCRISPR Optimal Target Finder (flycrispr.molbio.wisc.edu/tools). We targeted exon 1 of *D. simulans Gr32a* and *Ppk25*, and exon 2 of *Gr33a*. CRISPR oligos were annealed and ligated to plasmid pDCC6 {Addgene # 59985, (Gokcezade et al., 2014)} following restriction digest with *BbsI*. Sequences used to synthesize CRISPR oligos are provided in *Supplementary Table 1*. Plasmids were injected at 100 ng/uL concentrations for each of 2 - 3 plasmids targeting a single gene. Animals were screened for mutations by PCR followed by 15% non-denaturing PAGE (Zhu et al., 2014) or directly by sequencing. Please see *Supplementary Table 3* for details on results of CRISPR injections for *D. simulans*. All newly generated mutant strains were backcrossed at least 5 times to WT *D. simulans* before testing for behavior. Subsequent to this out-crossing to WT *D. simulans*, we mated heterozygous flies to obtain homozygous stocks for each allele. Given the absence of balancers in *D. simulans*, we verified genotypes at each generation by PCR analysis as described above to generate homozygous stocks.

### Generating *D. simulans* and *D. melanogaster* transgenic animals

To make Gr32a-GAL4 lines, we amplified the ∼3.8 kb region upstream of the Gr32a start codon from *D. simulans* or *D. melanogaster* (primer sequences provided in *Supplementary Table 1*) and subcloned it into pENTR/TOPO plasmid followed by Gateway-mediated subcloning into pBPGw. We then phiC31-integrated each DNA construct into Chr III landing sites for each species, sim986 for *D. simulans* and attp2 for *D. melanogaster* (Groth et al., 2004; Knapp et al., 2015; Stern et al., 2017). pJFRC2(10xUAS-ReaChR::Citrine) plasmid (Inagaki et al., 2014) was provided by David Anderson, and it was used to generate the Citrine reporter in *D. simulans* using the landing site described above. Embryo injections were performed by Rainbow Transgenics (Camarillo, CA) or BestGene (Chino Hills, CA).

### Molecular analysis of *Gr32a*, *Gr33a*, and *Ppk25* mutations in *D. simulans*

RNA was isolated from 10 WT or mutant *D. simulans* males (Trizol, ThermoFisher) and converted to cDNA using SuperScript III First-Strand Synthesis (Invitrogen, ThermoFisher). RT-PCR was performed using primers based on coding sequence (*Supplementary Table 1*) that spanned exon-intron junctions of the respective locus (*Gr32a*, *Gr33a*, or *Ppk25*) to avoid amplifying products from genomic DNA. Use of these primers did not generate detectable product in no-RT controls. We subcloned and sequenced RT-PCR products from flies mutant for each allele of *Gr32a*, *Gr33a*, and *Ppk25*; we also directly sequenced RT-PCR products from flies mutant for each allele of *Gr32a* (except *Gr32a*^*Δ26*^), *Gr33a*, and *Ppk25*. RNA isolation and the subsequent RT-PCR and sequencing were performed on 2-3 independent cohorts of WT and mutant flies. Sequence reads of subclones obtained from these RT-PCR studies and their alignment to the corresponding WT allele confirmed the presence of the expected mutation for each fly stock.

### Histology

Tarsi were dissected in ice-cold PBS, fixed in fresh 4% paraformaldehyde at 22°C, washed 3x in PBT, and then mounted as described before (Fan et al., 2013). Samples were imaged using a Zeiss LSM700 (Z-stacks) and processed in ImageJ.

### Tests for Non-Neutral Evolution

Alignments of genomes from 27 insect species (23 drosophilids, housefly, mosquito, honeybee, and beetle) were generated for coordinates (dm6: chr2L:11,110,412-11,114,209) encompassing the *D. melanogaster* Gr32a ∼3.8 kb regulatory sequence, and this alignment was subsequently downloaded from the Table Browser (UCSC Genome Browser, 2015 update) (Blanchette et al., 2004; Karolchik et al., 2004; Rosenbloom et al., 2014). PhyloP scores were computed for this region for three main tests: 1) a basewise “all-branches” test for conserved or accelerated evolution in all species compared to a neutral model (one test per nucleotide), 2) a whole-region “all-branches” test for conserved evolution in all species compared to a neutral model (one test for the whole region), and 3) a basewise “subtree” test for conserved or accelerated evolution in the designated species (*D. melanogaster* or *D. simulans*) compared to the other species (one test per nucleotide for each designate species) (Pollard et al., 2010). PhyloP scores are negative log_10_ P values of a likelihood ratio test comparing two evolutionary models (alternate vs. neutral or subtree vs. subtree complement). Scores near “0” indicate the expected rate of evolution, while large scores indicate conservation (phyloP score > 2) or acceleration (phyloP score < −2). PhyloP scores were tallied across coding sequence, introns, UTRs, and intergenic regions (Siepel et al., 2005). The phylogenetic model for neutral evolution was based on 4-fold degenerate sites in the 27-species genomic alignment and also downloaded from the UCSC Genome Browser. PhyloP scores and R code are made available for reproducible workflow in *Supplementary Folder 1* (Allaire et al., 2017; Team, 2017; Xie, 2016) (https://cran.r-project.org/doc/FAQ/R-FAQ.html#Citing-R).

### Hydrophobicity plot

Hydrophobicity scores were generated with ProtScale (Artimo et al., 2012) (<u>web.expasy.org/protscale</u>) using *D. melanogaster* or *D. simulans* Gr32a amino acid sequences as input. We used the Kyte and Doolittle hydrophobicity scale with a window size of 19 amino acids and uniform weights across all residues. The seven transmembrane domains were identified using HMMTOP (Tusnady and Simon, 1998, 2001) (www.enzim.hu/hmmtop) to predict the topology of Gr32a for both *D. melanogaster* and *D. simulans*.

### Courtship assays

All courtship assays were performed at zeitgeber time 6-10 at 22°C, illuminated by a fluorescent ring lamp (22W) suspended 4 cm above the courtship chamber and recorded with a Sony camcorder (HDR-XR550V) (Fan et al., 2013). Experiments performed under dark conditions were illuminated by red LEDs and recorded as above in a dark room. Virgin flies were collected at eclosion and light entrained (12 hours L/D, 25C) for 5-7 days prior to testing. Experimental males were kept in isolation and tested with flies that were group-housed (∼20 flies per vial) by species and sex. Foreleg tarsi were surgically removed at eclosion and males were tested as described above. We used w^501^ *D. simulans* as targets in male-male assays to distinguish them by eye color from test males. Behavioral assays were scored blind to genotype, using the MATLAB software ScoreVideo (Wu et al., 2009). We scored courtship as the period of time male flies spent chasing the stimulus fly, performing unilateral wing extension (courtship song), licking, abdominal bending (attempted copulation), or copulation. Courtship Index (CI) was calculated as the time spent by the male performing these behaviors, divided by the total assay time (15 minutes).

### Taste assay

Preference assays were performed as described previously (Montell, 2009). Briefly, 60-well plates were prepared the day prior to experimentation and kept at 4°C. Dyes were diluted from stock solutions (Brilliant blue FCF and Sulforhodamine B, 12.5 mg/ml each) and resuspended in agarose, to which sucrose or sucrose spiked with quinine-HCl were subsequently added. Final concentrations were: agarose (1%), Brilliant blue FCF (0.125 mg/mL; Wako Pure Chemical), Sulforhodamine B (0.125 mg/mL; SigmaAldrich), sucrose (1 mM; JT Baker), and sucrose (5 mM) spiked with quinine (0.5 mM; SigmaAldrich). Substrate with sucrose or sucrose spiked with quinine were randomly colored blue or red and counterbalanced for all experiments. 3-4 day old male and female flies were flipped into fresh food for 2 days at 12-hour light/dark cycle at 25°C. Flies were then food deprived by flipping them into vials containing 1% agarose and placed in the dark for 24 hours. Flies were then briefly anesthetized with CO_2_ and loaded onto the 60-well plates (zeitgeber time 2-3), which were put in a box and placed in the dark at 25°C for 90 min. Abdomens were scored as blue, red, purple (mixed eating), or no food coloring blind to genotype and color condition. A Preference Index was calculated for each 60-well plate as follows: (N^B^ + 0.5*N^P^)/(N^B^ + N^R^ + 0.5*N^P^) or (N^R^ + 0.5*N^P^)/(N^B^ + N^R^ + N^P^) where N^B^, N^R^, and N^P^ = total # flies with blue, red, and purple abdomens, respectively. Each genotype was tested ≥ 6 times.

### Statistical Analyses for Behavioral Data

We used Fisher’s exact test to analyze categorical data (e.g. percent assays with CI > 0.05) and we used the Bonferroni correction for multiple group comparisons as necessary. For other comparisons, we first tested whether data were normally distributed using a Lillefors’ goodness-of-fit test using MATLAB. Data not violating this assumption were analyzed with parametric tests (Student’s t-test for two groups or one-way ANOVA); otherwise, data were tested with a non-parametric test (Kolmogorov-Smirnov test for two groups or Kruskal-Wallis test). A Tukey’s post hoc test following multiple group comparisons was used to determine which groups differed significantly.

## AUTHOR CONTRIBUTIONS

O.M.A. and N.M.S. designed the fly experiments. O.M.A., K.M.T., P.H.S., and J.P. conducted the fly experiments and analyzed behavioral data. A.A.H. and K.S.P. conducted bioinformatics analyses of Gr32a ∼3.8 kb regulatory region. S.P. helped with molecular analysis of Gr33a and Ppk25 mutants. G.W.D. provided invaluable advice, resources, and laboratory space for some of these experiments. J.M.K. and D.L.S. provided reagents to generate transgenic *D. simulans* lines. O.M.A., A.A.H., K.S.P., and N.M.S. wrote the paper.

## ACKNOWLEDGEMENTS

We thank Drs. Thomas Clandinin, Pu Fan, Liqun Luo, and Devanand Manoli, members of the Shah Lab, and Z. Yan Wang for helpful comments during the course of this work or on the manuscript. We thank Dr. Anderson for sharing the pJFRC2[UAS-ReaChR::Citrine] plasmid. This work was performed in part under the auspices of the U.S. Department of Energy by Lawrence Livermore National Laboratory under Contract DE-AC52-07NA27344 (A.A.H.). It was funded by the National Science Foundation Graduate Research Fellowships Program (O.M.A.), Human Frontier Science Program Postdoctoral Fellowship (S.P.), Gladstone Institutes (A.A.H., K.S.P.), flexible funds from Career Awards in Biomedical Sciences from the Burroughs Wellcome Fund, Ellison Medical Foundation, McKnight Foundation for Neuroscience, and Sloan Foundation (N.M.S.).

## SUPPLEMENTAL FIGURE LEGENDS

**Figure 1–figure supplement 1:**
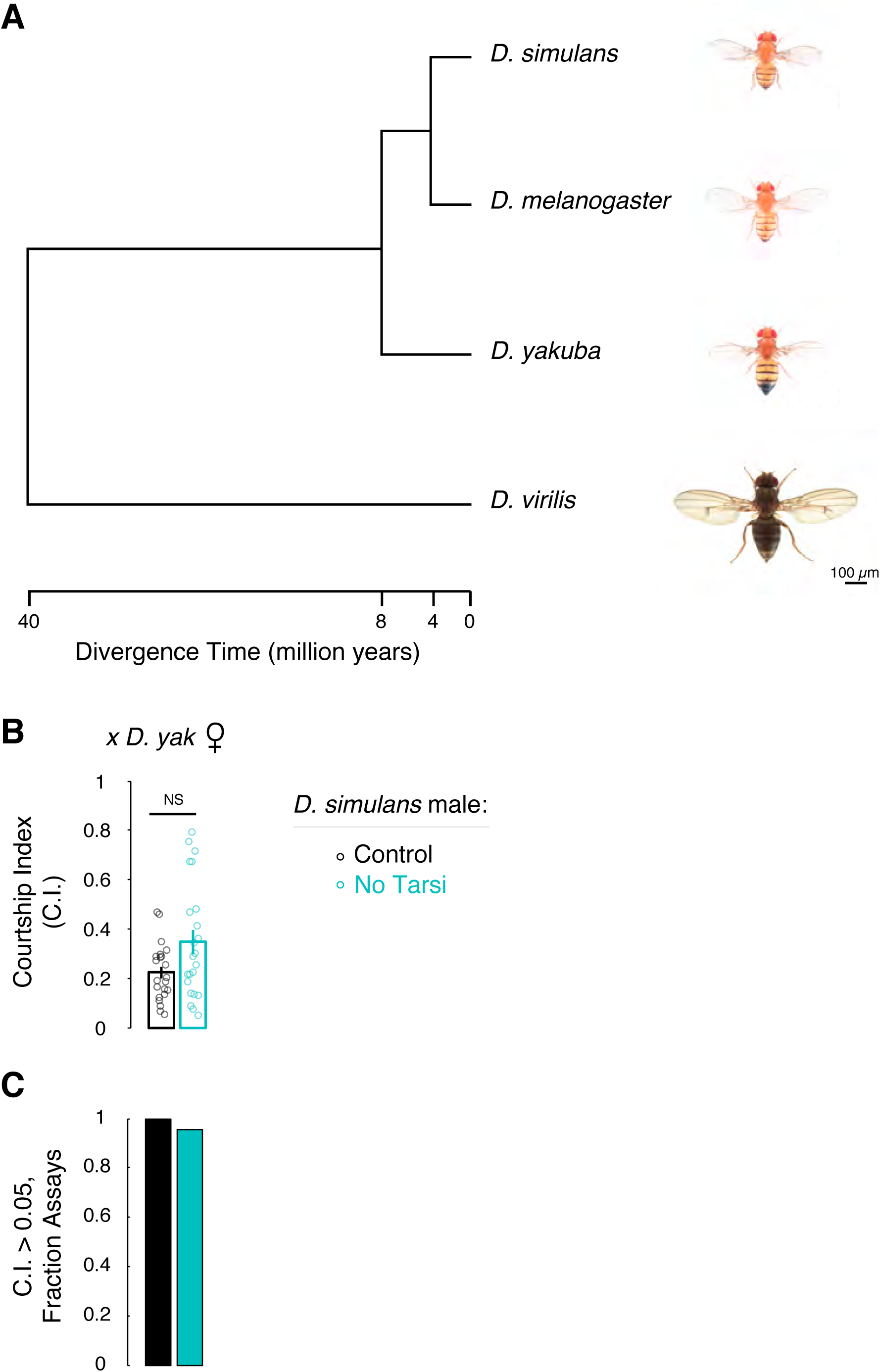
*D. simulans* foreleg tarsi inhibit courtship toward other drosophilids. (A) Evolutionary relationship of the four *Drosophila* species used in this study. Females of each species shown to scale and with representative pigmentation pattern. Images of these flies were obtained from FlyBase (Gramates et al., 2017). (B,C) Foreleg tarsi do not inhibit *D. simulans* males from courting *D. yakuba* females. Mean ± SEM; each circle denotes CI of a *D. simulans* male; n = 22 - 23/cohort; ***p<0.001.

**Figure 2–figure supplement 1:**
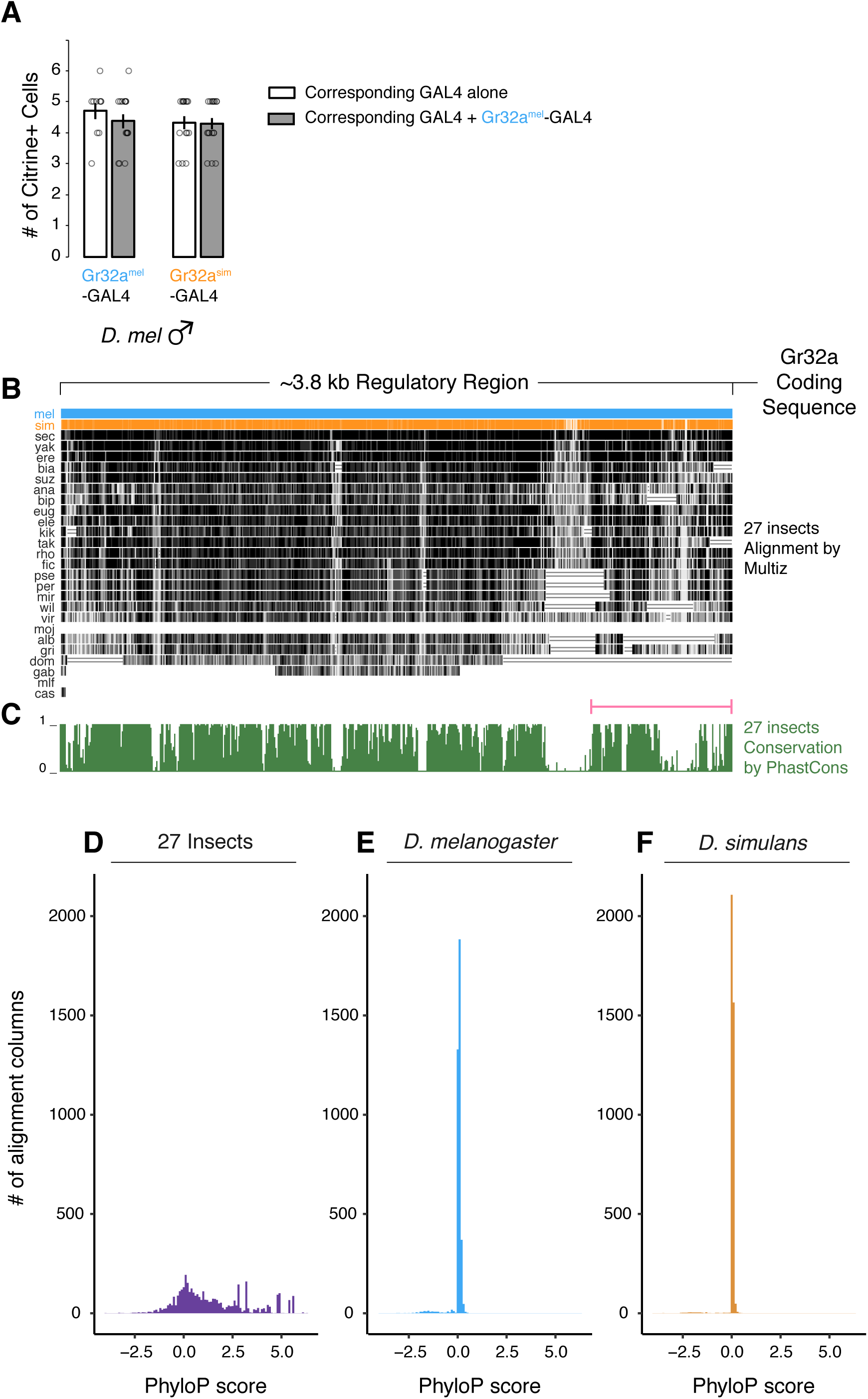
A regulatory region upstream of *Gr32a* coding sequence is conserved across drosophilids. (A) No difference in the number of Citrine+ cells in T4 and T5 foreleg segments of *D. melanogaster* males observed with either Gr32a^mel^-GAL4 or Gr32a^sim^-GAL4 alone or in combination. These findings indicate that the upstream regulatory sequence in Gr32a is functionally conserved between *D. melanogaster* and *D. simulans*; however, it is formally possible that the similarity in number of Citrine+ cells in *D. melanogaster* carrying one or both *GAL4* alleles reflects effects of transvection in the presence of both *GAL4* alleles rather than functional conservation. Mean ± SEM; each circle denotes Citrine+ cell count for a foreleg tarsum; n = 10 - 17/genotype. (B) 27-insect alignment of the ∼3.8 kb DNA element that drives *Gr32a* expression in *D. melanogaster* and *D. simulans*. mel, *D. melanogaster* (blue); sim, *D. simulans* (orange); sec, *D. sechellia*; yak, *D. yakuba*; ere, *D. erecta*; bia, *D. biarmipes*; suz, *D. suzukii*; ana, *D. ananassae*; bip, *D. bipectinata*; eug, *D. eugracilis*; ele, *D. elegans*; kik, *D. kikkawai*; tak, *D. takahashii*; rho, *D. rhopaloa*; fic, *D. ficusphila*; pse, *D. pseudoobscura*; per, *D. persimilis*; mir, *D. miranda*; wil, *D. willistoni*; vir, *D. virilis*; moj, *D. mojavensis*; alb, *D. albomicans*; gri, *D. grimshawi*; dom, *Musca domestica*; gab, *Anopheles gambiae*; mlf, *Apis mellifera*; cas, *Tribolium castaneum*. (C) Track showing PhastCons conservation score across the region in (B). Pink bar indicates the intergenic region directly 5’ of the Gr32a start codon and 5’UTR, which contains several blocks of highly conserved sequence. Higher peaks indicate higher likelihood of bases being in a strongly conserved element. (D) Distribution of nucleotide resolution phyloP conservation scores for the region shown in (C). Most bases in the region are evolving at the same or slower rate than 4-fold degenerate (4D) sites in the multiple sequence alignment of 27 insects. Large positive (> 2) or negative (< −2) scores indicate conservation or acceleration, respectively. Scores near “0” indicate a similar substitution rate to 4D sites. (E, F) Distribution of phyloP scores for branch-specific evolutionary tests. Most bases in the ∼3.8 kb region are likely evolving more slowly or as slowly as expected in *D. melanogaster* (E) and *D. simulans* (F) compared to the other 26 insects in the tree. Scores near “0” indicate a similar rate of DNA evolution in the designated species relative to all other species in the tree.

**Figure 3–figure supplement 1:**
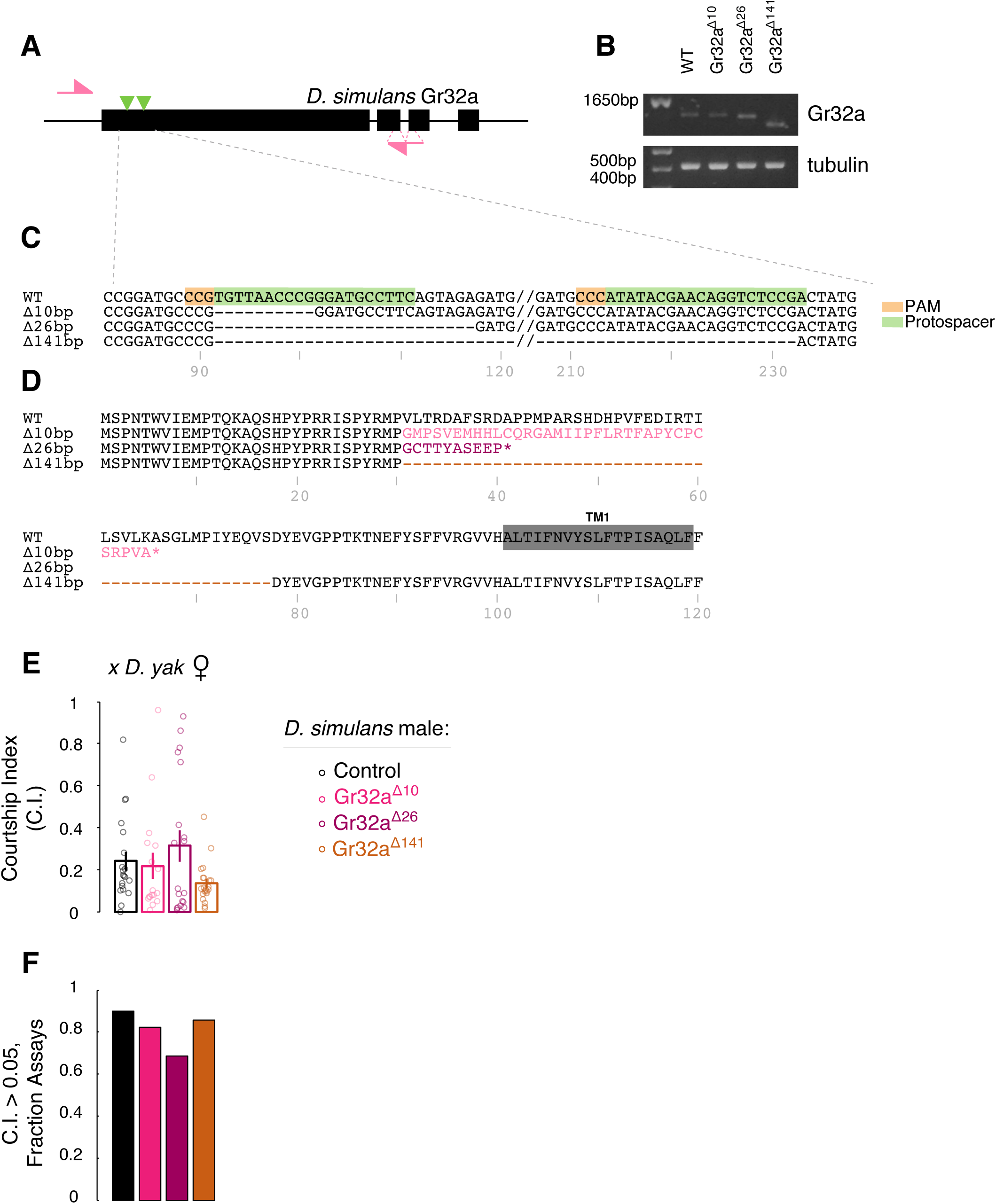
Generating Gr32a mutant *D. simulans* via CRISPR/Cas9. (A) Schematic of *D. simulans Gr32a* locus. Pink arrows, PCR primers; green triangles, CRISPR target sites; black rectangles, exons. (B) RT-PCR products for Gr32a and tubulin in WT and *Gr32a* mutant *D. simulans*, using PCR primers shown in (A). DNA ladder shown in first lane. (C) DNA sequence comparison of WT and mutant *Gr32a* alleles. PAM, Protospacer Adjacent Motif. (D) Predicted amino acid sequence of WT and mutant *D. simulans* Gr32a. The predicted first transmembrane domain (TM1) is highlighted in gray in the WT protein. *, premature stop codon. (E, F) No difference in courtship of *D. yakuba* females by WT and *Gr32a* mutant *D. simulans* males. Mean ± SEM; each circle represents CI of a male; n = 17 - 21/genotype (E, F).

**Figure 3–figure supplement 2:**
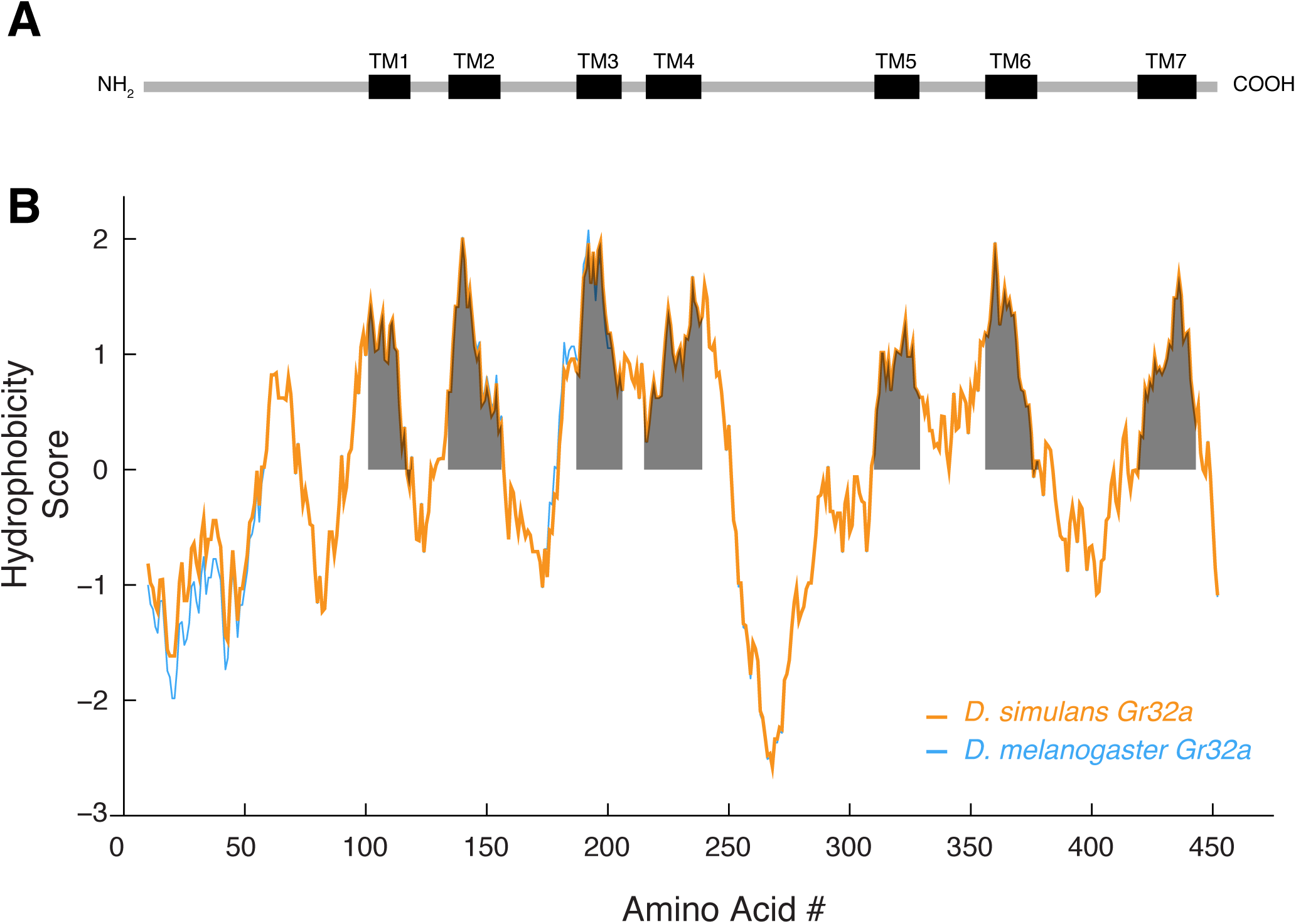
Hydrophobicity plot for Gr32a. (A) Predicted location of the seven transmembrane domains (black rectangles) in Gr32a based on plot shown in (B). The NH_2_ terminal is predicted to be intracellular. (B) Hydrophobicity plot of *D. simulans* and *D. melanogaster* Gr32a. Predicted transmembrane domains are shown by gray shading. Please see *Supplementary Table 2*.

**Figure 3–figure supplement 3:**
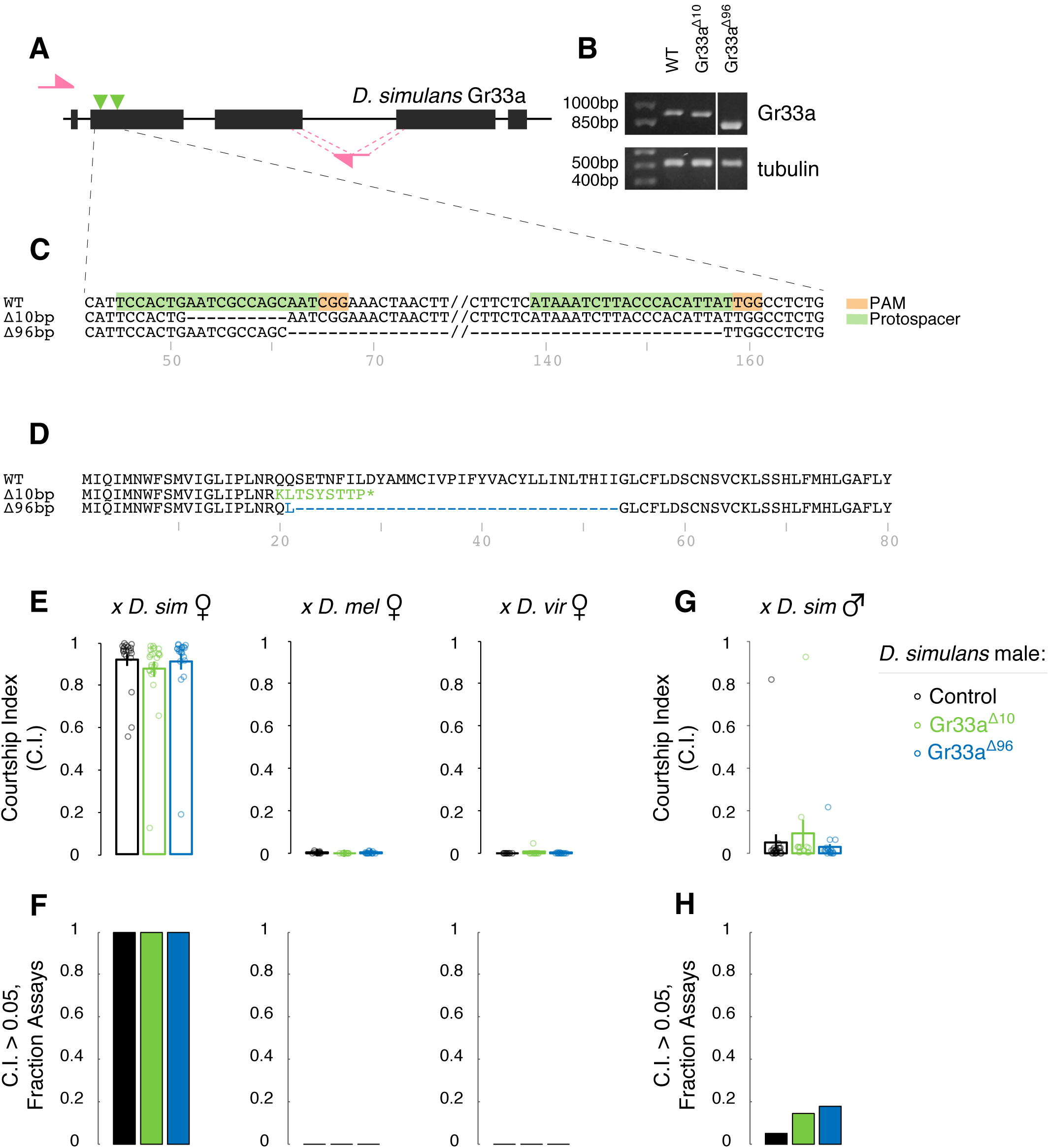
Gr33a is not required to inhibit interspecies courtship behavior of *D. simulans* males. (A) Schematic of *D. simulans Gr33a* locus. Pink arrows, PCR primers; green triangles, CRISPR target sites; black rectangles, exons. (B) RT-PCR products for Gr33a and tubulin in WT and *Gr33a* mutant *D. simulans*, using PCR primers shown in (A). Note that products from WT, *Gr33a*^*Δ10*^ and *Gr33a*^*Δ96*^ flies were run on the same gel, and lane between *Gr33a*^*Δ10*^ and *Gr33a*^*Δ96*^ has been cropped out for clarity of comparison. DNA ladder shown in first lane. (C) DNA sequence comparison of WT and mutant *Gr33a* alleles. Predicted amino acid sequence of WT and mutant *D. simulans* Gr33a. *, premature stop codon. (E, F) WT and *Gr33a* mutant *D. simulans* males court conspecific but not *D. melanogaster* or *D. virilis* females. (G, H) WT and Gr33a mutant *D. simulans* males show similar low levels of courtship toward conspecific males. Mean ± SEM; each circle denotes CI of one male; n = 10 - 24/genotype.

**Figure 3–figure supplement 4:**
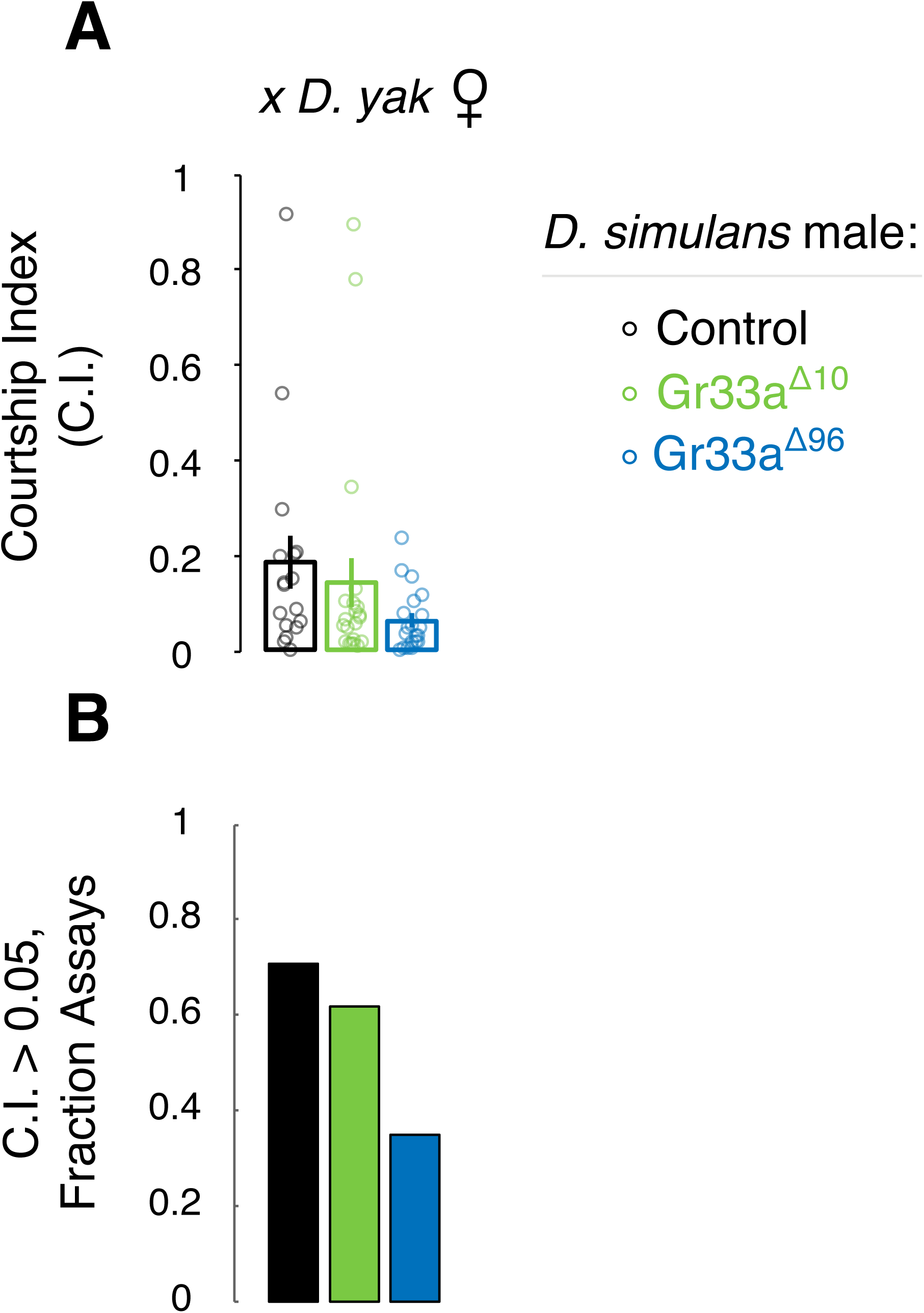
Gr33a does not regulate courtship toward *D. yakuba* females. (A, B) WT and *Gr33a* mutant *D. simulans* males court *D. yakuba* females at comparable levels. Mean ± SEM; each circle denotes CI for one male; n = 17 – 22/genotype.

**Figure 3–figure supplement 5:**
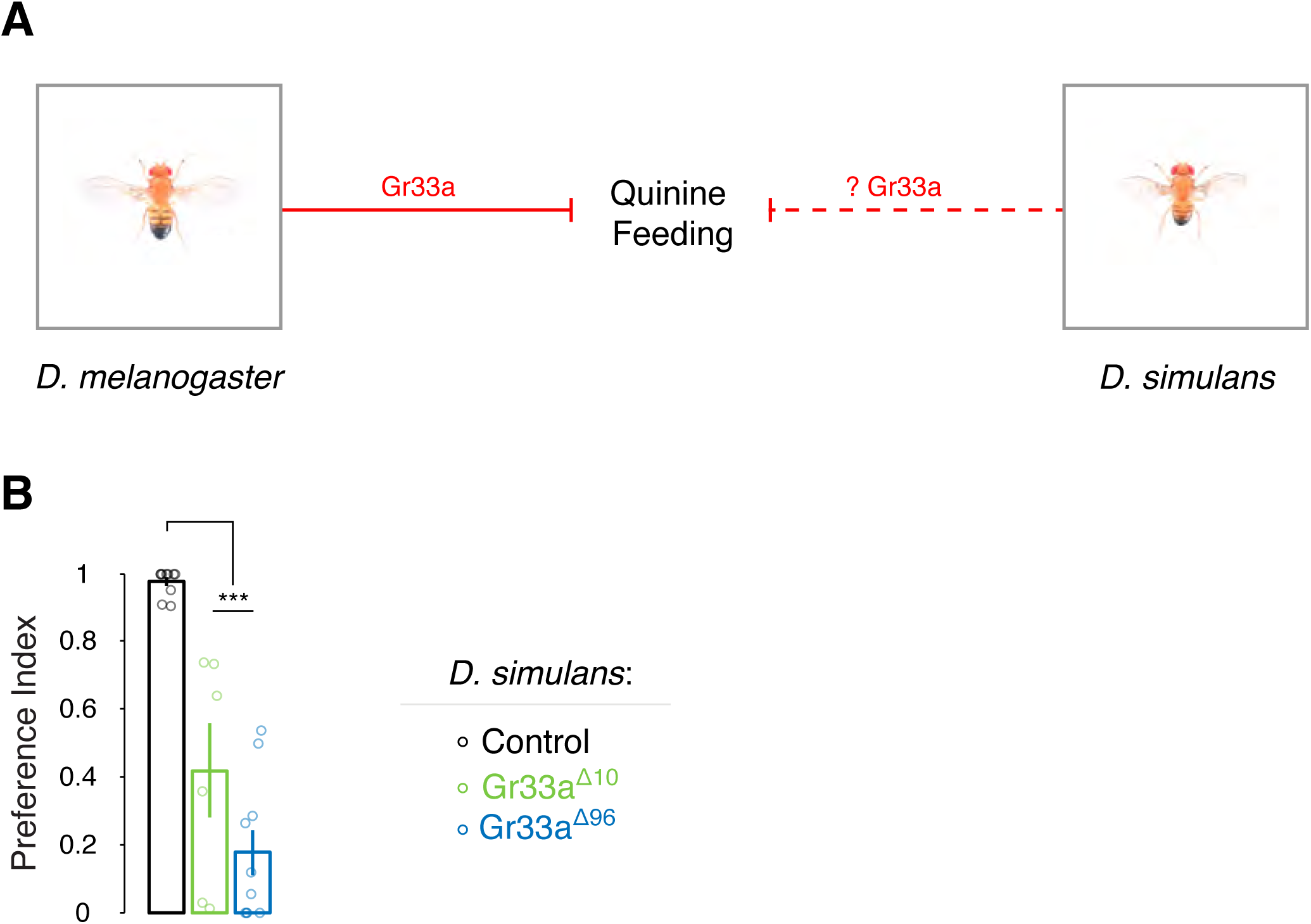
*Gr33a* inhibits *D. simulans* from feeding on quinine. (A) We tested whether, similar to *D. melanogaster*, Gr33a inhibits feeding on quinine-containing food in *D. simulans*. (B) Significant decrease in preference by *Gr33a* mutant *D. simulans* for food containing only sucrose. Mean ± SEM; each circle denotes Preference Index for one experiment; 90 ± 4 *D. simulans* of each genotype were used/experiment; n = 6 - 10 experiments/genotype; ***p<0.001.

**Figure 4–figure supplement 1:**
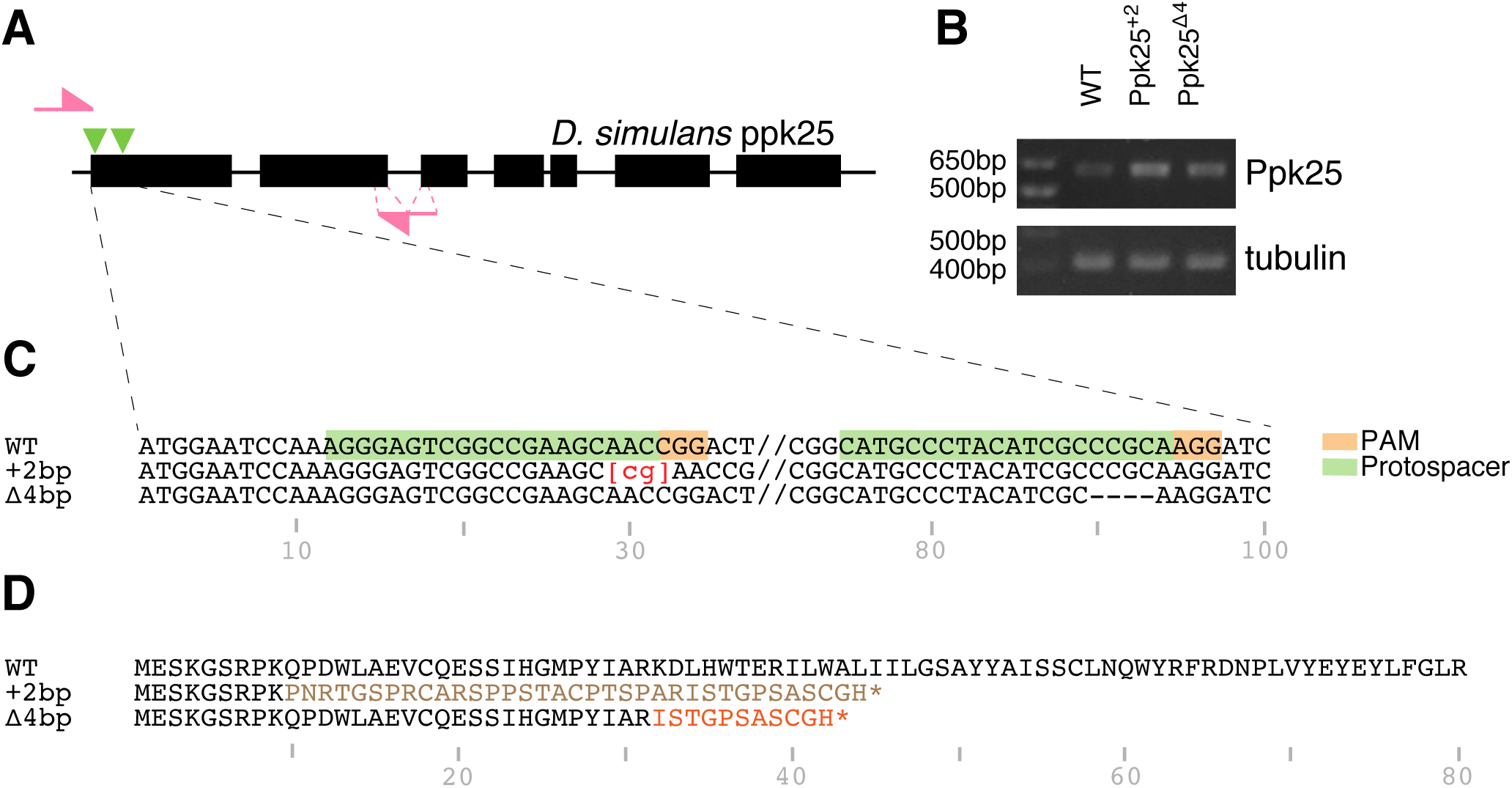
Generating Ppk25 mutant *D. simulans* via CRISPR/Cas9. (A) Schematic of *D. simulans Ppk25* locus. Pink arrows, PCR primers; green triangles, CRISPR target sites; black rectangles, exons. (B) RT-PCR products for Ppk25 and tubulin in WT and *Ppk25* mutant *D. simulans*, using PCR primers shown in (A). DNA ladder shown in first lane. (C) DNA sequence comparison of WT and mutant *Ppk25* alleles. (D) Predicted amino acid sequence of WT and mutant *D. simulans* Ppk25. *, premature stop codon.

**Figure 4–figure supplement 2:**
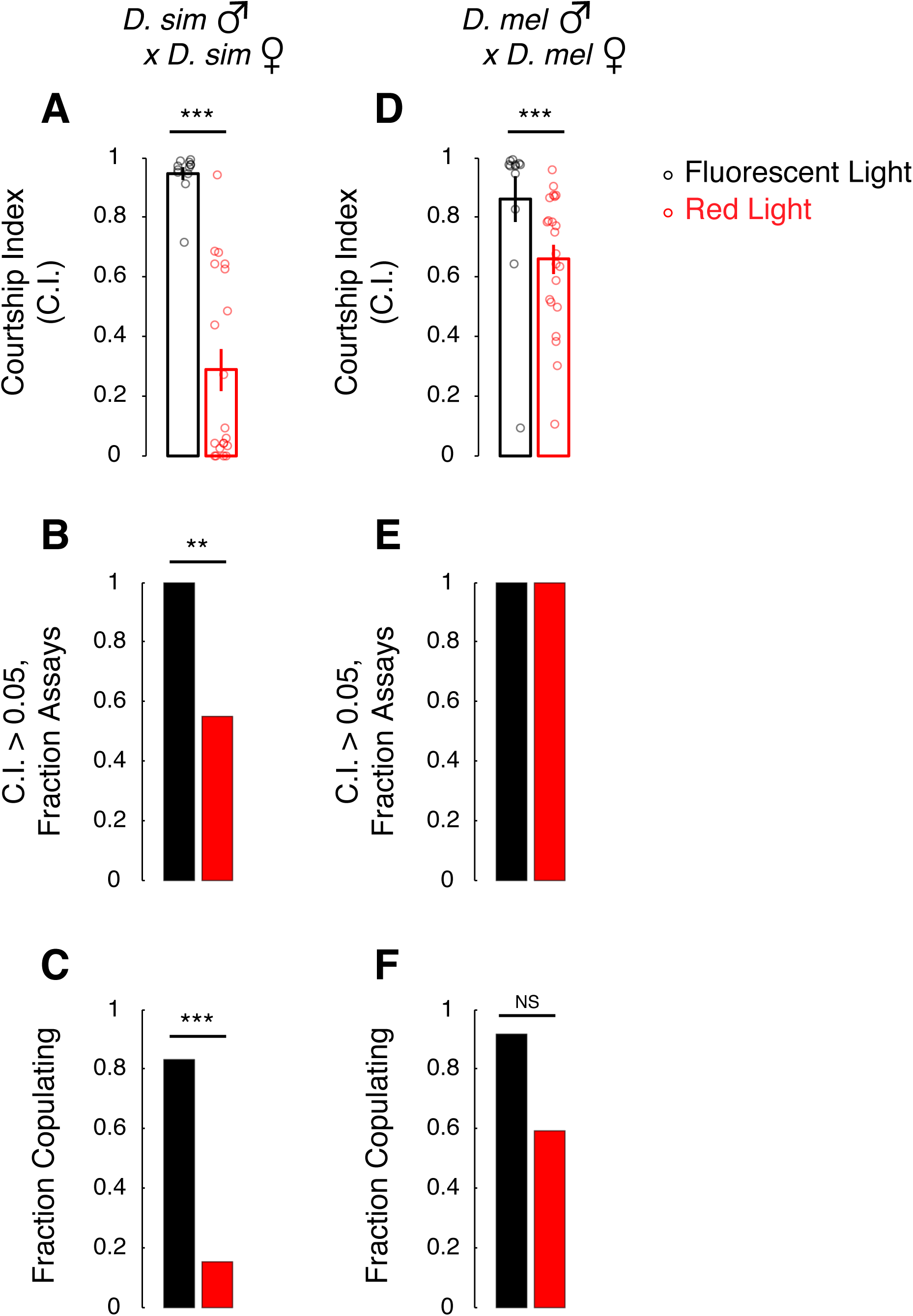
Light promotes conspecific courtship by *D. simulans* and *D. melanogaster* males. (A, D) *D. simulans* and *D. melanogaster* males court conspecific females less under red light-only illumination. (B, E) Fewer *D. simulans*, but not *D. melanogaster*, males court conspecific females intensely under red light-only illumination. (C, F) Fewer *D. simulans* males copulate under red light-only illumination whereas there is no difference in fraction *D. melanogaster* males copulating under red light-only or fluorescent illumination. Mean ± SEM; each circle denotes CI for one male; n = 12 - 22/cohort; **p<0.01; ***p<0.001.

**Figure 4–figure supplement 3:**
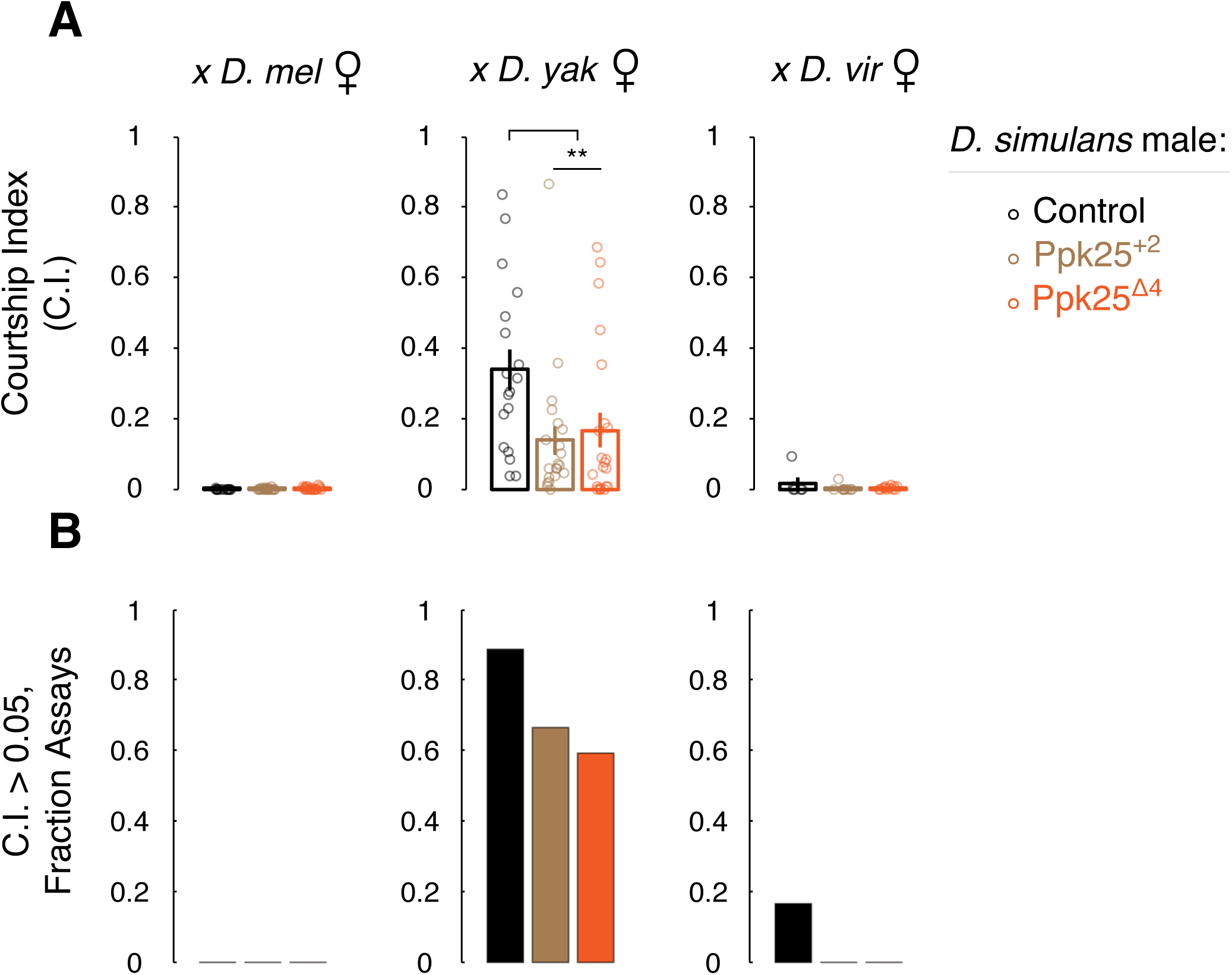
Ppk25 is not essential to inhibit interspecies courtship by *D. simulans* males. (A, B) No difference between *D. simulans* males WT or mutant for Ppk25 in courtship of *D. melanogaster* and *D. virilis* females. Mutant males court *D. yakuba* females less than WT (A). Mean ± SEM; each circle denotes CI for one male; n = 6-24/cohort; **p<0.01.

